# Interconnected Macroporous GelMA Scaffolds for Cell Culture through Temperature Controlled Spinodal Decomposition

**DOI:** 10.64898/2026.10.02.755893

**Authors:** Simon Dumas, David González Martínez, Adriene Peixoto De Oliveira, Daniel Ferri-Angulo, Lisa Bacchin, Alexis Arcas, Charline Blatché, Dominique Goudouneche, Raphaëlle Paounov, Marie Hut, Marwan Puaud, Jose Moran-Mirabal, Julie Foncy, Laurent Malaquin

**Affiliations:** LAAS-CNRS, 7 avenue du colonel Roche, Toulouse, France; McMaster University, Department of Chemistry and Chemical Biology, Hamilton, Ontario, Canada; CMEAB, Université Toulouse III Paul Sabatier, CMEAB, 133 route de Narbonne, Toulouse, France

## Abstract

Developing biomaterials that faithfully recapitulate the complex architecture of native tissues is central to tissue engineering. Conventional hydrogels typically exhibit nanoscale porosity, which limits mass transport and cell migration and thereby constrains tissue development. Here, we exploit liquid–liquid phase separation (LLPS) to generate interconnected macroporous gelatin methacryloyl (GelMA) scaffolds with tunable pore sizes ranging from 2 to 50 µm under physiological pH conditions. By mapping the GelMA/dextran phase diagram, we identify a composition range in which LLPS occurs through spinodal decomposition, resulting in a spontaneous bicontinuous, trabecular-like architecture. Pore size can be further tuned by varying polymer concentrations and temperature. We advantageously combined thermal gradients and photopatterning to enable spatial control over pore morphology to fabricate hierarchically structured materials. The biological relevance of these architectures is assessed using fibroblasts cultured within controlled porosity gradients. We show that pore size strongly influences cell infiltration, growth, and spatial organization, with larger pores providing sufficient space for cell migration while reducing the dependence on matrix degradation. Pore size of 7–15 µm is identified for maximizing fibroblast growth. Overall, our approach provides a versatile platform for porosity screening, enabling the engineering of biomimetic hydrogels with optimized microarchitectures tailored to specific biological contexts for three-dimensional cell culture.

## Introduction

One of the major challenges in tissue engineering is to reproduce the structural and functional complexity of the extracellular matrix (ECM). Native tissues are intrinsically heterogeneous: ECM architecture, density, porosity, stiffness, and permeability vary across space and over multiple length scales, defining local environments with distinct mechanical, biochemical, and transport properties.^1^ Similarly, cells are not uniformly distributed within this matrix but are organized into functional compartments known as cellular niches, whose density and interstitial space also vary considerably across tissues.^1^ Reproducing such heterogeneity is therefore essential for engineering matrices that more faithfully recapitulate native tissues. Importantly, cell behavior is not governed by a single material parameter, but by the combined influence of matrix mechanics, physical confinement, biochemical cues, and molecular transport, which together regulate cell fate, organization, and tissue maturation..^1–6^ Most current approaches rely on hydrogels,^7^ which can be either synthetic (e.g., PEGDA, PVA) or natural (e.g., collagen, chitosan, gelatin).^8^ While synthetic hydrogels offer high reproducibility, they often lack biodegradability and cannot be remodeled by cells. Among natural hydrogels, gelatin methacryloyl (GelMA) has attracted considerable interest because it combines the biological advantages of natural polymers with the mechanical tunability of synthetic systems while being biodegradable.^9^

However, whether natural or synthetic, hydrogels inherently exhibit porosity at the nanometric scale, which poses significant limitations.^10^ Their densely crosslinked polymer networks severely restrict the diffusion of oxygen, nutrients, and metabolic waste, which often leads to the formation of necrotic regions in constructs exceeding a few hundred micrometers in thickness. Furthermore, their dense structure limits cell migration, spreading, and endothelial infiltration, processes that are critical for vascularization and functional tissue integration. This is commonly addressed by using softer gels with a looser polymer network. However, this presents a trade-off between cell mobility and substrate rigidity. In most hydrogel-based materials, stiffness and porosity are intrinsically coupled: dense matrices support cell adhesion but restrict migration and mass transport, whereas highly porous gels are often too soft to maintain cell identity and material structural integrity. This trade-off hinders tissue maturation, underscoring the need for precise control over porosity scales in hydrogels. An ideal hydrogel would be stiff enough to support cell adhesion while remaining porous enough to allow cell mobility and mass transport.^1,11,12^

Several strategies have been proposed to address these limitations.^12,13^ Freeze-drying and ice templating locally disrupts the polymer chains, introducing macroporosity.^14–18^ Another approach involves the introduction of porogens before crosslinking. Foaming^19,20^ or emulsification^21^ have been employed, as well as the use of a solid sacrificial material such as gelatin, hyaluronic acid, calcium carbonate or alginate beads to create voids in the matrix.^22,23^ However, these methods often lack pore interconnectivity, as the pores are induced by discrete objects such as beads or bubbles. To overcome this limitation, granular materials were used as building blocks, where interstitial space between the blocks serves as the macroporosity.^24–30^ Other methods leverage additive manufacturing to create porous scaffolds in a bottom-up manner. Techniques such as electrospinning,^31^ extrusion bioprinting,^32–36^ and two-photon polymerization^37,38^ provide a high degree of spatial control over microarchitecture. Nevertheless, these methods are often time-consuming, difficult to scale up, and may suffer from limitations in either resolution or printing speed.

Recently, liquid-liquid phase separation (LLPS), also referred to as aqueous two-phase separation (ATPS), has emerged as a promising approach for introducing porosity in hydrogels.^39^ This approach employs an additional polymer (e.g., PEG, dextran, or PVA) that separates from the biomaterial prior to crosslinking, allowing for the introduction of macroporosity at the cell-scale (5-100 µm) in a single step. The primary challenge lies in controlling the phase separation kinetics to achieve the desired pore size and interconnectivity. In pioneering studies, the porosity was controlled by modifying the LLPS kinetics through fine-tuning of experimental parameters such as pH, polymers concentrations, or by the addition of viscosity enhancers.^40–43^ However, the pores dimensions mostly depend on gel formulation, and these methods do not allow for the LLPS control in space and time. More recently, Polymer-Induced Phase Separation (PIPS) has emerged as a highly adaptable method for controlling porosity. In this approach, pore size is determined by the progressive demixing of two polymers triggered by photopolymerization. PIPS is particularly powerful, as it enables precise tuning of porosity by simply adjusting the exposure dose, eliminating the need for fine adjustments of concentration and pH. However, since this method uses light to control both liquid-liquid phase separation (LLPS) and crosslinking, the porosity and mechanical properties remain intrinsically coupled and cannot be independently adjusted.^44–46^

Despite these advances, offering fine control of the hierarchical porosity with high interconnectivity and physiological pH is still a challenge, and there is a growing interest for lab-ready LLPS-based methods. Moreover, the specific parameters that drive the system toward either a globular morphology (resulting in discrete voids) or a spinodal, trabecular morphology (yielding interconnected structures) still require thorough characterization. In this study, we address some of these challenges. We first thrive into the LLPS process of GelMA/dextran systems at physiological pH in a wide range of concentrations and temperatures, providing guidelines to interpenetrated porous hydrogels. We propose a method to spatially shape the material and porosity at multiple scales by combining temperature gradients and photopatterning. Then, we demonstrate the applicability of the material for cell culture using fibroblasts seeded in the material with a wide range of porosities. We show that porosity significantly impacts cell growth in three dimensions, emphasizing the importance of controlling microarchitecture at multiple scales in tissue engineering.

## Results

### 1. Porous GelMA scaffolds generation through LLPS

We exploit LLPS to generate porous scaffolds using synthetic GelMA (type A, bloom 300, SD 75%) as a biomaterial and dextran (Mn = 500 kDa) as a porogen agent. GelMA/dextran solutions remain fully mixed at 37°C, while cooling below the GelMA gelation temperature (typically <20°C for porcine gelatin) triggers LLPS. Phase separation results from the self-assembly of gelatin molecules into collagen-like triple helices, which physically cross-links GelMA, modifying its water solubility relative to dextran (Figure 1A). Depending on the GelMA/dextran ratio, LLPS proceeds through either binodal or spinodal regimes. In the binodal regime, the system is metastable, and local fluctuations must overcome an energy barrier to trigger nucleation and droplet growth, resulting in a discrete dispersion of one phase in the other. In contrast, spinodal separation is thermodynamically favorable and proceeds spontaneously by diffusion without an energy barrier, generating an interconnected biphasic structure. This spinodal regime is particularly interesting in the context of biomaterials as it allows the cells to move freely within the tridimensional space, without the need for degrading the matrix, as well as facilitating interstitial flows for nutrient supply. However, spinodal decomposition occurs only in a narrow and highly specific thermodynamic regime, where the system is intrinsically unstable, corresponding to a limited region of the bicomponent system phase diagram.

**Figure 1:**
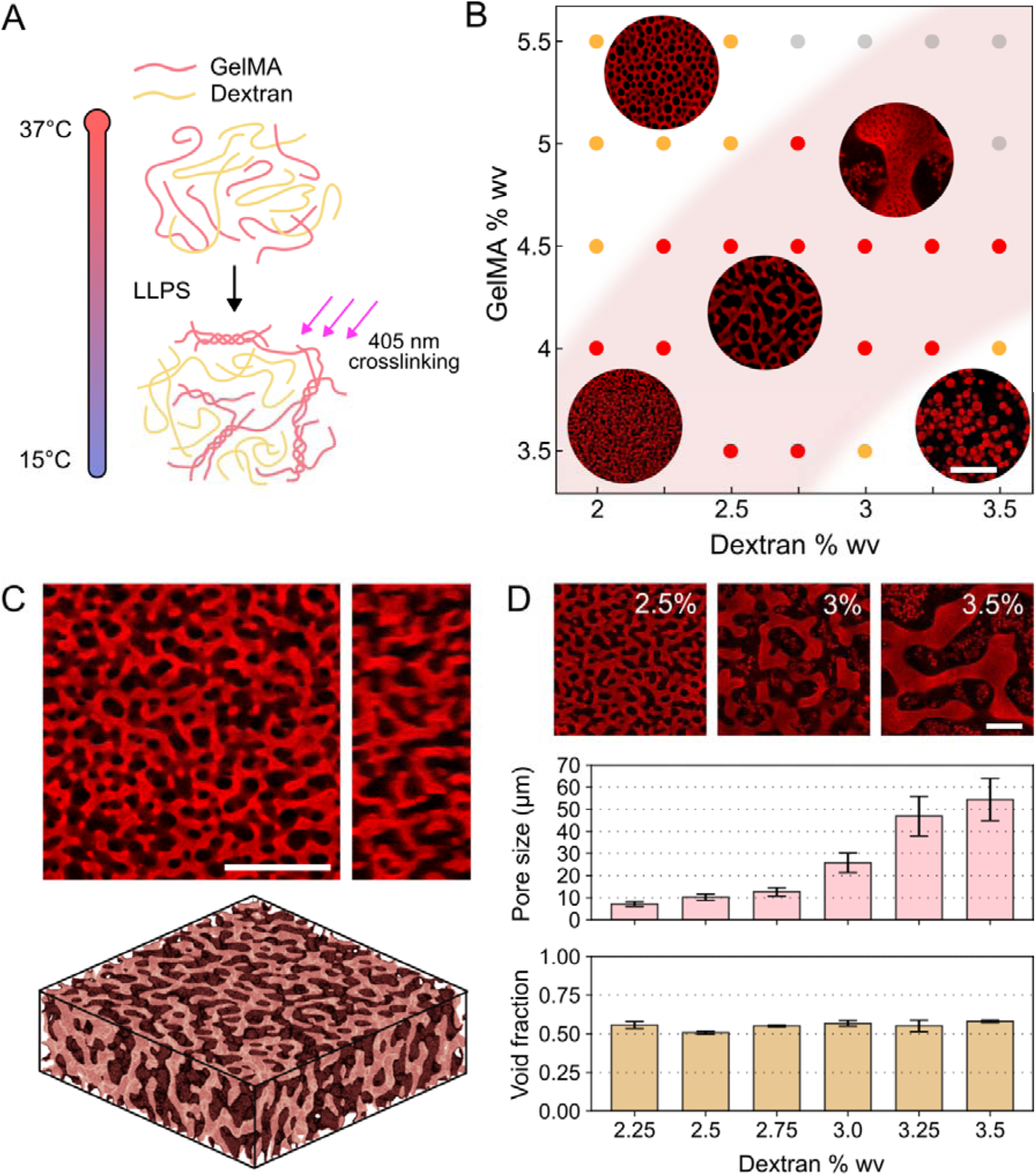
GelMA scaffolds generated through LLPS by cooling down to 15°C. **A.** Principle of the LLPS. **B.** Phase diagram of the GelMA/dextran (500 kDa) system. The pink background represents the spinodal region of the diagram. Yellow dots: binodal regime. Red dots: spinodal regime. Grey dots: fully demixed or not exploitable. **C.** Top: 3D confocal imaging of a LLPS-induced gel containing 4.5% GelMA 2.5% dextran. Bottom: 3D reconstruction of the same picture using Volume Viewer Fiji plugin. **D.** Influence of dextran concentration for a fixed GelMA concentration of 4.5% (n = 3 pictures in each sample). All scale bars 50 µm.

To place these different regimes in an experimental phase diagram, we performed LLPS on 35 solutions of GelMA/dextran in PBS with concentrations spanning from 3.5–5.5% w/v for GelMA and 2–3.5% w/v for dextran. We cooled down the samples to 15°C in a microfluidic chamber to induce LLPS and subsequently photopolymerized them to immobilize the phases in the biphasic configuration. We established a phase diagram and identified three regimes (Figure 1B). At high GelMA/low dextran concentrations ([GelMA]/[dextran]□2), the binodal regime produces dextran droplets within a GelMA matrix. Conversely, at low GelMA/high dextran concentrations ([GelMA]/[dextran]□1.1), GelMA droplets form within a dextran matrix (Figure 1B). In between these two regimes, we identified a band where LLPS occurs following the spinodal decomposition (Figure 1B, S1) and generates a three-dimensional interconnected porosity, as confirmed using confocal microscopy (Figure 1C).

In the spinodal regime, the porosity directly links with the concentration of both components, characterized by larger pores when increasing the concentration of either GelMA or dextran (Figure S1). The evolution of porosity is very sensitive to even small changes of concentration. For example, at a fixed GelMA concentration of 4.5%, increasing the dextran concentration from 2.25% to 3.5% enabled precise control over scaffold pore size, ranging from 7.1 to 53.3 µm (Figure 1D). Overall, simply tuning both polymer concentrations allowed us to modulate scaffold pore size across a wide range of approximately 2–50 µm (Figure S1), which is relevant to a broad variety of biological models, as it encompasses the size of many cell types as well as larger structures such as some blood and lymphatic vessels, and some cellular niches. Interestingly, the void fraction, defined by volume ratio of dextran to GelMA phases, remains stable around □_v_ ≈ 0.6 over all conditions (Figure 1D).

### 2. Nanoarchitecture and mechanical properties of LLPS gels

To further characterize the macroporous material and understand the relationship between the microarchitecture with the morphology at nanometer scale and the mechanical properties, we performed transmission electron microscopy (TEM) and rheometric analysis. We compared three conditions: a soft gel containing 4.5% (w/v) GelMA (G4.5), a stiff gel containing 9% (w/v) GelMA (G9) and an LLPS-induced gel containing 4.5% (w/v) GelMA and 2.5% (w/v) dextran (G4.5D2.5) similar to the one of Figure 1C. TEM images (Figure 2A) showed that at higher GelMA concentrations, polymer chains pack more densely, with an apparent solid fraction rising from □□= 0.29 (soft G4.5) to □□= 0.39 (stiff G9). In the LLPS-induced gel, the polymer condenses during phase separation, reaching a solid fraction of □□= 0.40 in the gel areas (Figure 2B). This behavior aligns with our expectation that, in a macroporous structure with a void fraction near 0.5, the gel phase must roughly double its concentration to fit in the phase volume. Consequently, the gel in G4.5D2.5 more closely resembles G9 than G4.5 but occupies only half of the volume.

**Figure 2:**
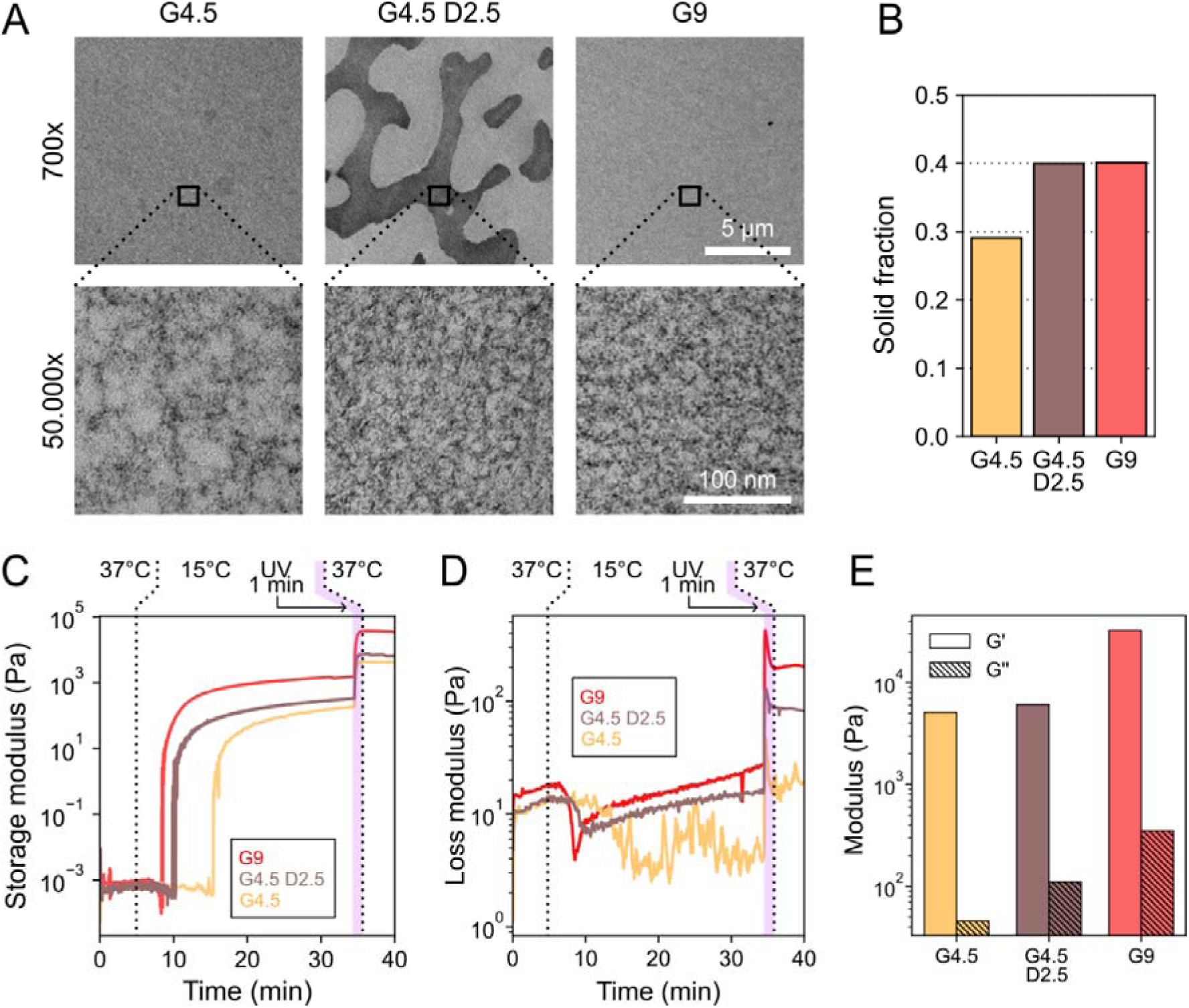
Structural and mechanical properties of LLPS-induced gels. **A.** TEM images of gels prepared at 15°C with three compositions: a soft gel (G4.5), a stiff gel (G9) and an LLPS-induced gel (G4.5D2.5). TEM images at 50.000x zoom allows us to see the polymer density at nanometer scale. **B.** Solid fractions observed in the 50.000x images for the three conditions. **C.** Storage modulus G’ upon time during phase separation and UV crosslinking. **D.** Loss modulus G’’ during the same sequence. **E.** Final values of G’ and G’’ in the three gels after LLPS, UV cross-linking and final equilibration to 37°C.

To assess the impact of macroporosity on mechanical properties, we performed rheological measurements on the same three conditions. Cooling the solutions to 15°C increased the storage modulus G’ by several orders of magnitude across all conditions, indicating physical cross-linking of GelMA (Figure 2C), while the loss modulus G’’ remained largely unchanged (Figure 2D). After 30 min at 15°C, UV irradiation was applied, leading to a significant increase in both G’ and G’’ (Figure 2E). After UV cross-linking and equilibration to 37°C, we observed G’ >> G’’ in all conditions, indicating that all cross-linked gels behave predominantly as elastic solids (Figure 2E). When comparing soft (G4.5), stiff (G9), and LLPS-induced (G4.5D2.5) samples, we found that G’ depends primarily on the total GelMA concentration rather than microarchitecture: it increased from 5.1 to 6.1 kPa upon dextran addition to G4.5, whereas it reached 32.6 kPa in the stiff (G9) gel. Furthermore, G’’ increased in the LLPS-induced gel, suggesting that the microporous architecture and dextran presence introduce an additional viscous component.

Overall, the mechanical properties of the material are primarily governed by the total GelMA content even if the gel densifies locally during LLPS. However, bulk-scale rheological measurements do not fully capture the local mechanical environment experienced by cells. To address this limitation, atomic force microscopy (AFM) measurements at the microscale will be essential in the future to correlate bulk properties with the actual rigidity perceived at the cellular scale in LLPS-induced hydrogels.

### 3. Dynamics of LLPS with temperature

To investigate how temperature influences the LLPS process, we developed an integrated platform to control the temperature in real time while imaging under microscope (Figure 3A). The system is based on Peltier thermoelectric elements with PID feedback control to precisely heat or cool a coverslip with a sub-0.1°C temperature accuracy. The device functioning and fabrication are described in Figure S2. Using this system, we were able to observe the phase separation process in real time under microscope (Figure 3B, Video S1). To quantify the phase separation dynamics over time, we developed an FFT-based image analysis pipeline that determines, in real time, the characteristic length of the main periodic features on the image. This approach enables pore size measurement directly using a bright field microscope, while yielding results consistent with confocal imaging (Figure S3). By displaying the image’s mean power spectrum over time in a kymograph, we can track the emergence of porosity as a peak appears, with its vertical position in the kymograph being inversely proportional to the pore size (Figure 3C).

**Figure 3:**
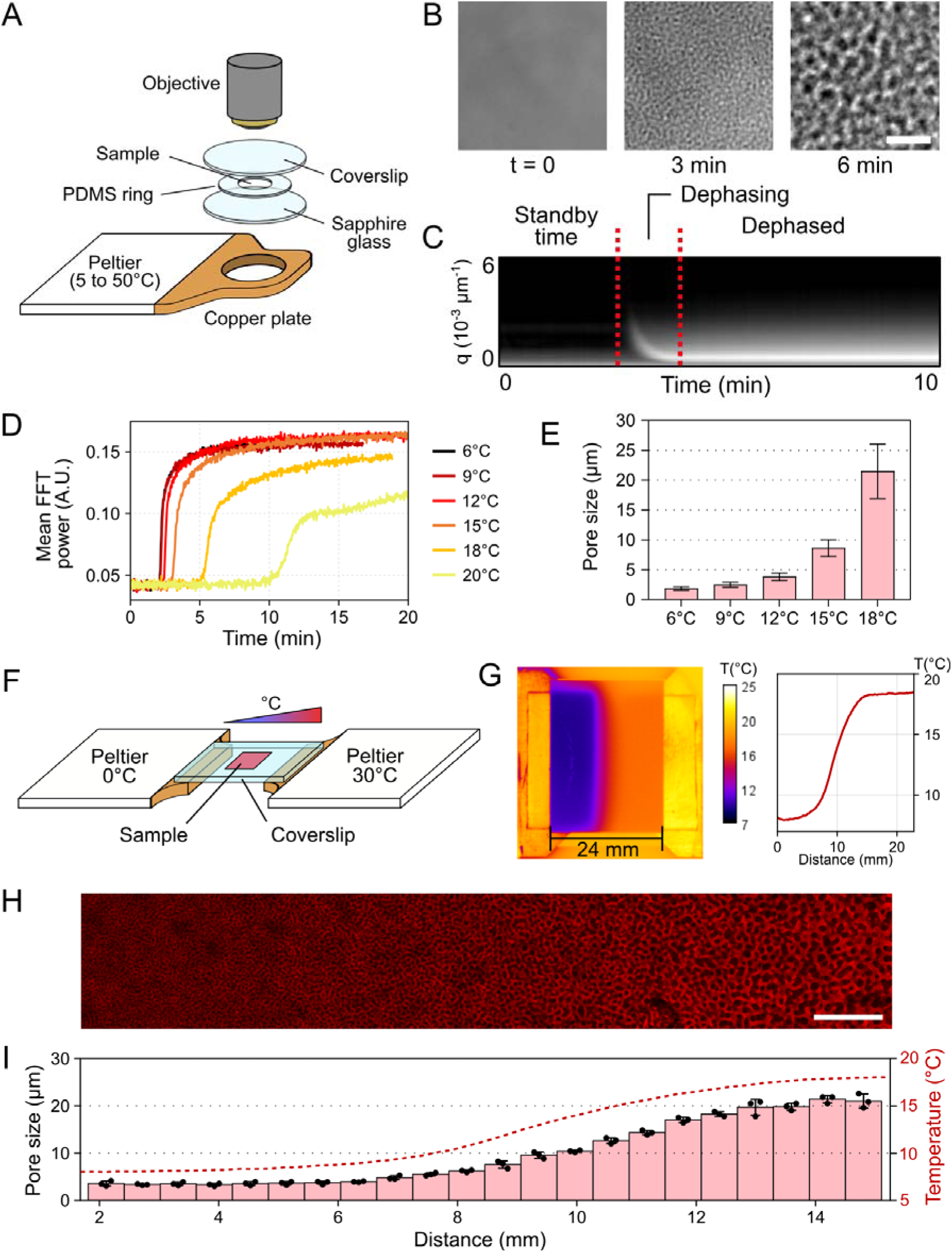
Dynamics of the spinodal decomposition with temperature. **A.** Schematic of the temperature control device. A programmable Peltier element controls the temperature of a copper plate with a precision of <0.1°C. The copper plate presents a hole on which is placed a high thermal conductivity sapphire glass window. The sample is pipetted between the sapphire window and a coverslip, separated with a PDMS ring (200 µm thick). The whole device is mounted on a microscope for real-time imaging of the LLPS process (see Figure S2 for more details). **B.** Phase contrast images of the gel during LLPS (18°C). **C.** Kymograph showing the evolution upon time of the FFT power spectrum for the 15°C condition. The vertical axis represents the spatial frequencies of features (in px^-1^), inversely proportional to the feature size (in px). Thus, a bright peak getting closer to the bottom with time indicates the formation and growth of pores in the gel. All kymographs are presented in Figure S4. **D.** Mean power spectrum over time for LLPS temperatures in the 6-20°C range. **E.** Final pore size for all temperature conditions. **F.** Schematic of the double-Peltier device. **G.** Thermal infrared pictures showing the temperature gradient (8-19°C) established on a glass coverslip. **H.** Gradient of porosity observed through confocal microscopy, fabricated using a temperature gradient of 8-19°C and a G4.5D2.5 solution. **I.** Porosity measured using the FFT method (Figure S2) on gels obtained using a temperature gradient of 8-19°C. All scale bars 50 µm.

Using this approach, we analyzed the phase separation dynamics in a 4.5% GelMA 2.5% dextran solution (G4.5D2.5) across a temperature range of 6–20°C (Video S2). FFT kymographs obtained at each temperature are presented in Figure S4 and are recapitulated in Figure 3D showing the evolution of mean FFT power over time. The increase in mean FFT power indicates textural changes in the images, marked by the emergence of a peak in the FFT spectrum. These results demonstrate that phase separation occurs within minutes at low temperatures (<10°C). As temperature increases, the phase separation is delayed, starting after 5 min at 18°C and 10 min at 20°C. This temperature dependence reflects the competition between GelMA chain association upon cooling and thermal agitation, which maintains molecular separation. At temperatures ≥21°C, no phase separation was observed, even after 30 min. This is in agreement to the fact that LLPS is triggered when physical gelation occurs, as GelMA remains liquid at temperatures above 21°C in our range of concentrations (1-10% w/v).

Importantly, porosity increases with LLPS temperature (Figure 3E), with pore sizes ranging from 1.8 to 21.5 µm across the 6–18°C temperature range while using the same solution. This trend arises from the balance between phase separation dynamics and gelation kinetics: at higher temperatures, phase separation accelerates due to increased chain mobility, while gelation proceeds more slowly, allowing additional time for phase growth. Importantly, the LLPS remained in the spinodal regime for all temperature conditions, demonstrating that the regime of phase separation mainly depends on the gel composition rather than the phase separation dynamics. This result is important because it allows us to tune the porosity in-situ without having to modify the gel precursor solution.

We leveraged this concept to generate controlled porosity gradients. The Peltier-based platform was enhanced by adding a second cooling element placed side by side with the first (Figure 3F). When a coverslip is placed, centered and in contact with both elements, a linear temperature gradient forms across its surface, as confirmed by infrared imaging, with the left and right Peltier elements set to 8°C and 19°C, respectively (Figure 3G). Using the same sample as before (G4.5D2.5), we generated a linear porosity gradient ranging from 4 to 23 µm over a 5 mm distance (Figure 3HI). This approach enables one-step fabrication of GelMA scaffolds with macroporosity gradients, thereby providing a versatile experimental platform for systematic investigation of porosity effects on cellular behavior within a single construct.

### 4. Photopatterning of LLPS-induced gels

The LLPS is fully reversible: heating the gel back to 37°C can dissolve non-photocrosslinked material (Figure 4A, Video S3). We exploited this feature to demonstrate the ability of photopatterning to fabricate various porous GelMA-based objects. For photopatterning, we employed a digital micromirror device (DMD)-based system (Primo, Alvéole) to expose patterns onto the samples mounted on the Peltier cooling system. To optimize the process, we applied varying light intensities to a tree-shaped pattern featuring structures of different width (Figure 4B). Our observations revealed that at low irradiation doses (<6 mJ/mm²), the structures were highly deformable and prone to significant swelling, resulting in poorly defined thin features. Conversely, at high doses (>10 mJ/mm²), small structures were well-defined, but undesired crosslinking occurred between branches.

**Figure 4:**
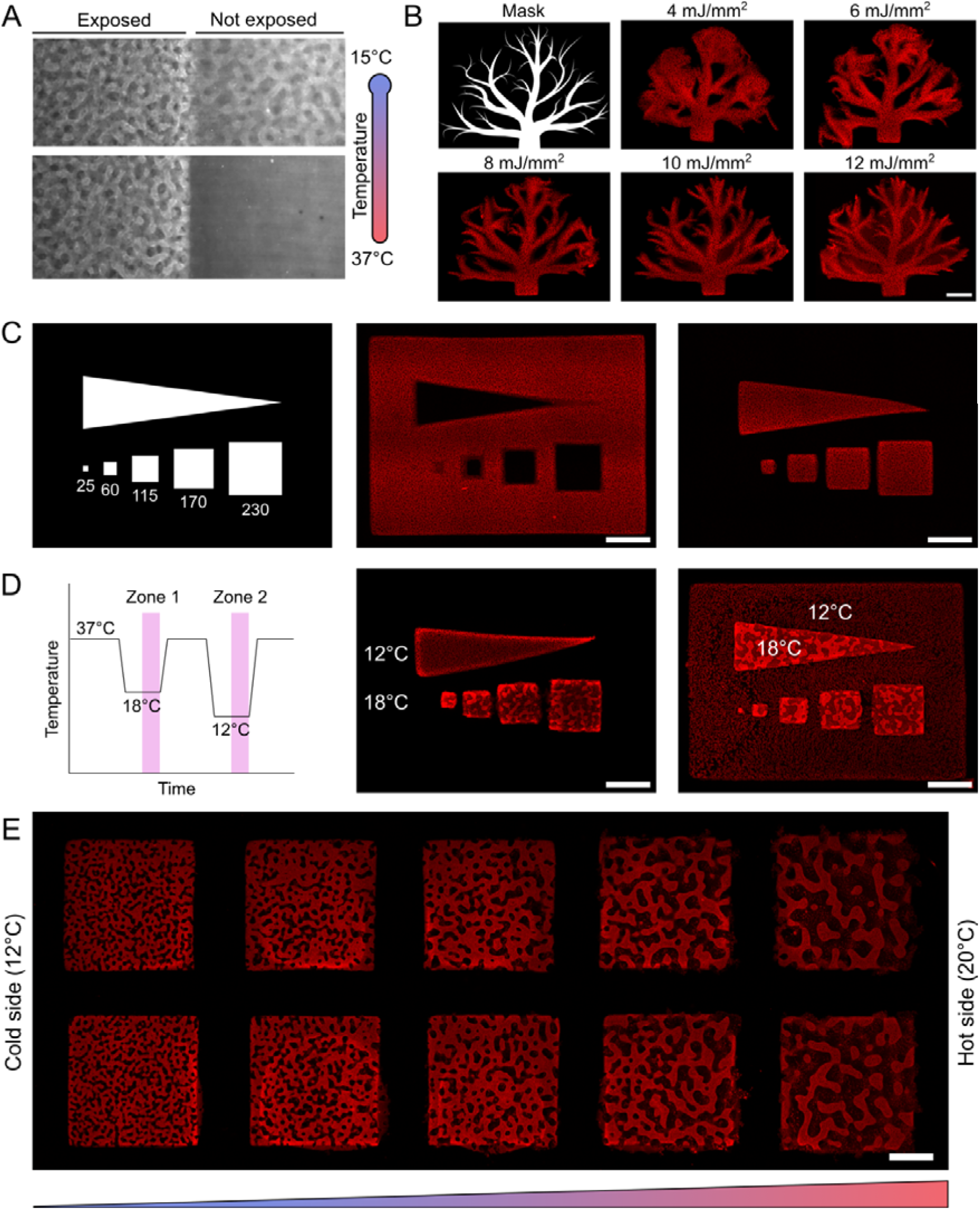
Photopatterning of LLPS-induced gels. **A.** Spinodal decomposition is reversible, the structures can dissolve when placed at 37°C unless photopolymerized (Video S2). **B.** Tree-shaped patterns printed on macroporous GelMA at different UV doses. **C.** Resolution test using either negative (middle image) or positive (right image) polarity. **D.** Multi-porosity printing using temperature cycles. **E.** Photopatterning within a temperature gradient to generate multi-porosity patterns. All scale bars 200 µm.

This phenomenon is inherent to the polymerization process itself: radicals generated by the photoinitiator (LAP) diffuse during illumination, and inhibitory oxygen in the immediate vicinity of the patterns is consumed, preventing it from fulfilling its inhibitory role over prolonged exposure times. This limitation may also arise from increased light scattering due to the presence of the two phases. Then, resolution tests demonstrated that both positive (isolated gel shapes) and negative (holes inside the shapes) structures could be fabricated (Figure 4C). However, achieving negative polarity proved more challenging due to the reasons described above: oxygen is consumed more rapidly while radicals diffuse and are less renewed within the confined geometry of a closed shape compared to an open one. Consequently, we were unable to produce holes smaller than ≈100 µm without internal photopolymerization. For positive polarity however, the feature size was less constrained, and small structures of 60 µm were successfully printed. However, very small objects (e.g. 25 µm squares), although well polymerized, frequently detached from the substrate due to the limited adhesion area.

By exploiting both the reversibility of LLPS and the temperature-dependent control of pore size, we can generate multi-porosity patterns on the same substrate by applying successive dephasing cycles at different temperatures (Figure 4D). This allows for the creation of either spatially distinct regions with varying porosities or nested structures with hierarchical porosity (Figure 4D). Additionally, multi-porosity objects can be obtained by combining photopatterning with LLPS under a temperature gradient to fabricate objects with spatially evolving porosity in a single step (Figure 4E).

Together, these approaches introduce a novel strategy for multi-scale GelMA structuring, where LLPS enables fine-tuning of the architecture at the cell scale (1–20 µm), while photopatterning defines larger features (>100 µm). These combined functionalities open new avenues for fabricating microphysiological systems with precise architectural control across multiple scales.

### 5. Macroporous gels enhance cellular development

To demonstrate the applicability of LLPS-induced gels for 3D cell culture, we seeded human lung fibroblasts (HLFs) within the gel. Unlike some studies where LLPS is induced by acidification^42,43^, our method operates under physiological pH and ionic conditions in PBS, enabling direct cell seeding into the gel precursor solution before phase separation. To evaluate cell viability and spreading, we cast gels containing 700,000 cells/mL using a PDMS chip as a mold on methacrylated glass coverslips and photopolymerized them after an 8-minute incubation at 15°C or 18°C to trigger LLPS. The gels were prepared with either no porosity (G9), small pores (∼7 µm, G4.5D2.5 at 15°C), or large pores (∼20 µm, G4.5D2.5 at 18°C) (Figure 5A). After 14 days in culture, the results demonstrated that porosity significantly enhanced cell proliferation. In non-porous gels, cells remained round and exhibited minimal motility, whereas in porous gels, cells elongated and proliferated. Cells rapidly invaded the porous matrix, increasing the occupied volume approximately 15-fold over 14 days, while it remained constant in non-porous gels (Figure 5B). These results demonstrate that porosity significantly improves cell mobility and spreading, and strong calcein signal confirms cell viability over two weeks. Confocal imaging also confirmed that the cells grow in three dimensions and are evenly distributed over the volume (Figure 5CD, Video S4).

**Figure 5:**
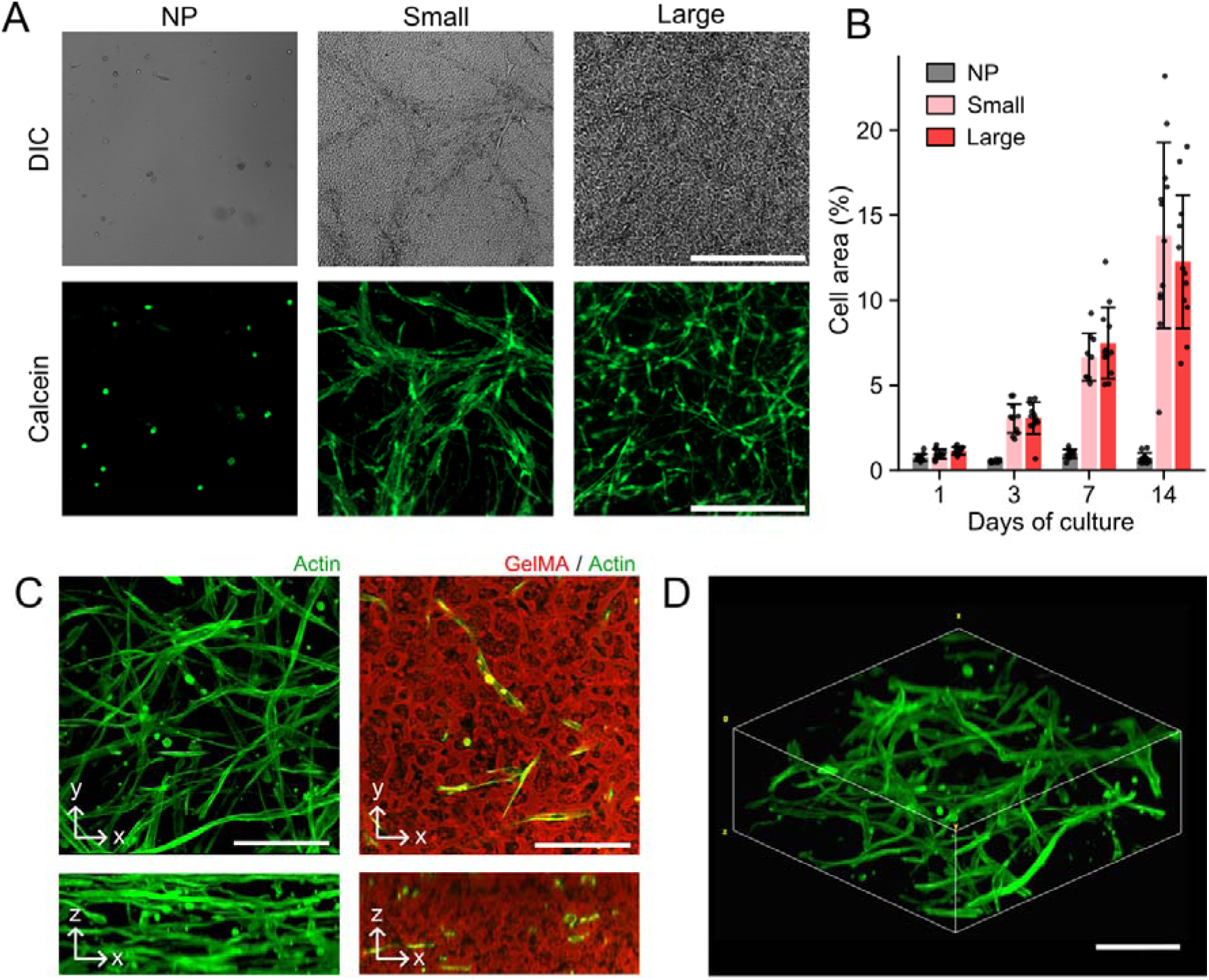
HLF growth on LLPS-induced gels. **A.** Cells observed after 14 days in three gels: NP (G9), Small (G4.5D2.5, 15°C) and Large (G4.5D2.5, 18°C). Gels were casted between two coverslips with a PDMS spacer ring of 200 µm thickness, then placed at either 15°C (Small) or 18°C (Large) for 8 min before photo-crosslinking. Scale bars 500 µm. **B.** Apparent area occupied by cells in the gel at different stages of the culture (n=12 pictures for each sample). **C.** Confocal imaging of the cells. Left: Maximum projection of actin channel; Right: a middle plane showing both actin and GelMA channels. Scale bars 100 µm. **D.** 3D reconstruction of actin channel in the same stack. Scale bar 100 µm.

### 6. Impact of porosity on cellular fibroblasts development

To study the impact of porosity on cell development, we seeded cells in a gel comprising a porosity gradient as in Figure 3HI. To produce this gel, we cast the gel precursor solution (G4.5D2.5) on methacrylated glass coverslips using a PDMS microfluidic chip as a mold. The chip was subjected to a temperature gradient (8-19°C) using the double-Peltier device as in Figure 3FG. After LLPS and photopolymerization, the PDMS chip was removed and the molded gel constructs containing cells remained attached to the glass coverslip and were cultured for 15 days (Figure 6A). The pores gradually increased from ≈ 4 to ≈ 23 µm within a 7 mm distance (Figure 6B). After 15 days in culture, this experiment showed that cell development is enhanced with increasing pore size, with a higher cell density in the larger pore regions (Figure 6CD). By counting the number of nuclei in each region, we can correlate the number of cells with porosity (Figure 6EF). These results demonstrated that cells proliferated more rapidly in regions with larger pores: cell numbers increased approximately 6-fold in regions with >10 µm pores after 7 days of culture, while only 2-fold in regions with <7 µm pores (Figure 6E). Interestingly, we obtained a maximum of proliferation with pores between 7 and 15 µm (Figure 6F), while porosities higher than that did not increase the proliferation rate. This result is in accordance to previous results^44^ where it was shown that increasing pore size does not systematically improve cell growth, which could be a consequence of reduced surface for the cells to adhere in a same volume, and optimal porosity seems to approximately match nuclei size (Figure S6-AB). This also shows the interest of using gradients to optimizematerial properties, as suggested in other studies^33,47,48^. Moreover, we observed a morphology change in response to pore size: cells in larger porosity regions (15-20 µm) grew homogeneously while the ones in the more confined regions (2-4 µm) arrange into bundles (Figure S6-AC). This may be due to lack of space, forcing the cells to remodel the matrix. This aligns with confocal microscopy observations of the hydrogel, where we observed cells digging galleries in low-porosity regions (2-7 µm), whereas in areas with larger pores, the cells adhered to the pore walls without modifying the scaffold microarchitecture (Figure 6G). Overall, these observations suggest a progressive shift in migration behavior with increasing confinement, from predominantly amoeboid migration in highly porous regions to mesenchymal and ultimately collective migration in more confined environments.

**Figure 6:**
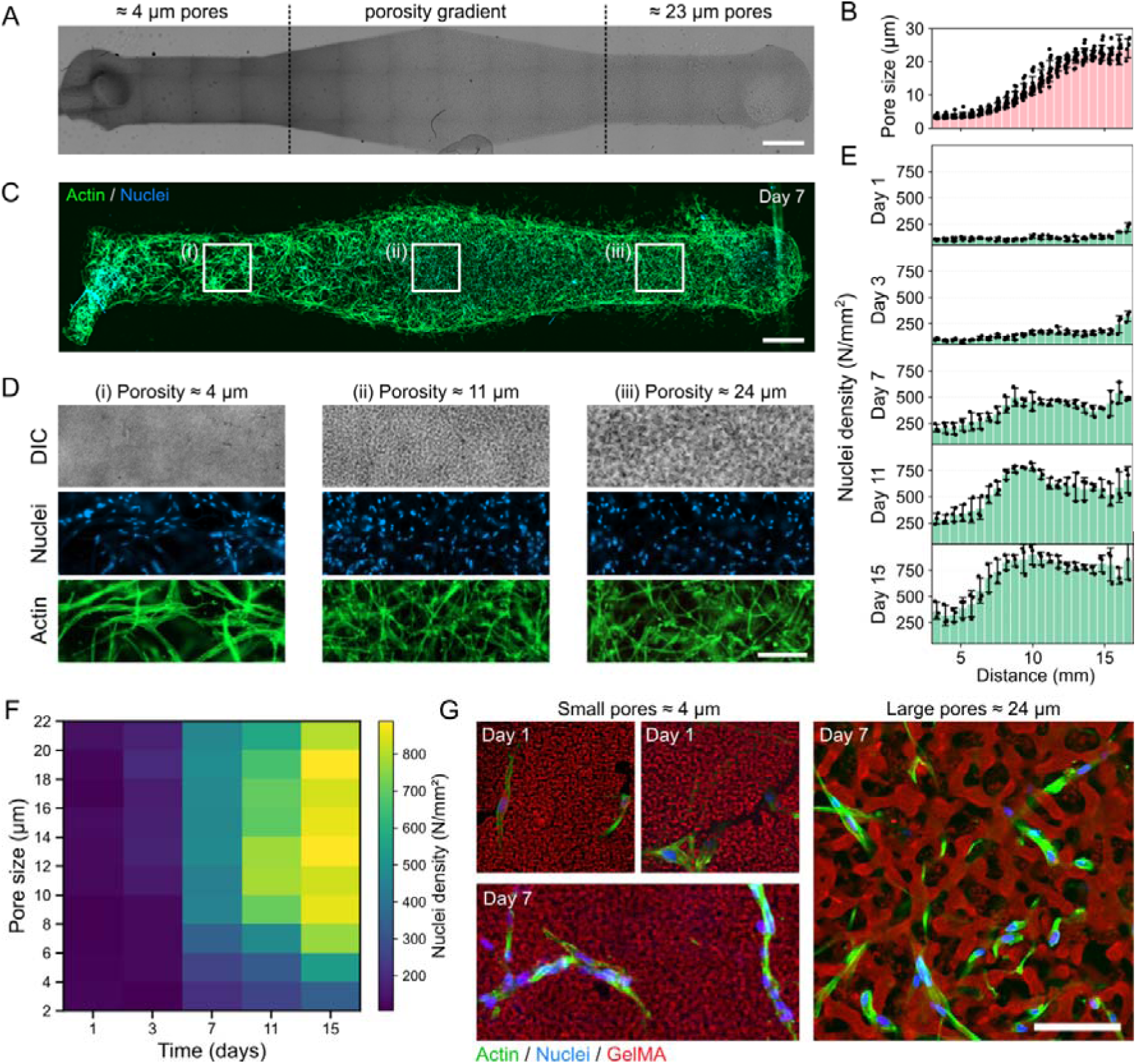
HLF culture in a gradient of porosity. **A.** LLPS-induced gel cast on a glass coverslip. LLPS was performed in a gradient of temperature (8-19°C), resulting in a porosity gradient with pores ranging from 4 to 23 µm. Scale bar 1 mm. **B.** Porosity over the distance in the gels (n = 15 samples). **C.** Cells grown in a porosity gradient gel after 7 days of culture. Scale bar 1 mm. **D.** Zoom in three regions of the same image with increasing porosity (i) 4 µm (ii) 11 µm, (iii) 24 µm. More timepoints are shown in Figure S5. Scale bar 200 µm. **E.** Number of nuclei in the gels divided into 30 sections with increasing porosity (n = 3 samples per day). **F.** Nuclei density relative to pore size at 1, 3, 7, 11 and 15 days of culture in the gel. **G.** Confocal images at day 1 and 7 with gels of small (4-6 µm) pores and at day 7 for large (20 µm) pores demonstrating an active matrix remodeling in confined environment. Scale bar 100 µm.

## Discussion

The development of biomaterials with tunable properties that promote cell growth is central in the field of tissue engineering. Conventional approaches rely on empirical optimization of the hydrogel formulation, where rigidity and porosity are inherently coupled, making it impossible to independently tune these critical parameters. This often results in compromised performance, as materials either lack mechanical stability or restrict cell mobility. To tackle this challenge, the field is evolving towards materials incorporating voids or channels.^1,39,49^ Notably, introducing porosity at the cell-scale is particularly desirable as it enables cell mobility without requiring extensive matrix degradation, allowing them to expand freely while maintaining a phenotype closer to their natural state. Porous scaffolds support rapid 3D colonization in just a few days, thus promoting tissue self-organization. Additionally, interconnected porous networks is known to enhance fluid flow, improving nutrient and oxygen delivery and reduces the risk of necrosis commonly observed in dense systems like spheroids.

Our method exploits LLPS to create GelMA scaffolds with a interconnected network of pores. We provide an thorough characterization of the LLPS kinetics in the spinodal regime as a function of hydrogel composition and temperature, establishing the mechanistic basis of pore formation. This understanding supports a predictable and reproducible fabrication method, capable of generating porous scaffolds with a wide range of pore sizes (2–50 µm) at scales relevant for diverse tissue engineering applications. Experimentally, the process can be performed in a straightforward manner by cooling down the sample in a mold before photopolymerization, but can also be coupled with photopatterning to produce more complex systems with multiple compartments and spatially organized porosities. By photopatterning in microfluidic devices, this feature can benefit to the organ-on-chip community to create complex systems. Most importantly, our results with fibroblasts culture demonstrate that cells develop more efficiently in macroporous environment, confirming the potential of our method to provide a more physiologically relevant environment for tissue development.

Despite these advantages, some challenges remain. Cooling the gel may stress temperature-sensitive cells, potentially compromising their viability, particularly for fragile cell types such as primary or stem cells. Nevertheless, we did not observe any significant impact on cell viability in this study, and the cooling step can be limited to less than 5 minutes at temperatures between 10 and 15°C, which should be compatible with most cell culture protocols. Another limitation arises from the cooling process itself: the sample cools from the outside inward, leading to temporary temperature gradients that can induce convective flows and result in an inhomogeneous phase separation. We were able to successfully obtain homogeneous samples up to 500 µm in thickness, but fabricating larger objects may be more challenging. While scaling up the fabrication by molding may present difficulties, the process can be compatible with extrusion bioprinting using a temperature-controlled nozzle and printing plate, or using FRESH bioprinting^47^, which will be explored in future studies.

Compared to other LLPS-based methods, our approach holds some improvements. Several macroporous bioinks have recently been developed and proved enhanced cell growth and tissue maturation. However, many of these materials are based on binodal (nucleation and growth) phase separation which do not offer the same level of control over microarchitecture in terms of pore dimensions and interconnectivity. A few studies have exploited spinodal decomposition on GelMA to generate interconnected porosity,^39–41^ and our method offers several improvements, most notably the ability to induce phase separation under physiological conditions (pH and PBS), enabling direct cell encapsulation, where previous studies have relied on acidic conditions (pH 4.1–5) to trigger phase separation and control pore size.^40,41^ More recently, Polymerisation Induced Phase Separation (PIPS) has gained attention for creating interconnected gels with controllable pore dimensions with cells embedded in the gel.^42–45^ In PIPS, phase separation is triggered by light during photopolymerization, with pore size determined by the balance between polymerization kinetics and diffusion of the polymer chains. A major advantage of PIPS is its compatibility with large-scale processing and volumetric printing, as it eliminates the need for temperature control. However, since light drives both phase separation and network cross-linking, mechanical properties and porosity remain intrinsically coupled, limiting independent control over these parameters. In contrast, our approach uses two distinct mechanisms: temperature first sets the porosity, and light then tunes the rigidity. This decoupling allows for independent adjustment of porosity and mechanical properties, offering a unique advantage to study the interplay between cellular confinement and matrix stiffness.

Moreover, beyond structural benefits, the material is well adapted to microphysiological systems. By combining photopatterning with LLPS, we can create complex, multi-property systems or organ-on-chip devices with a high degree of spatial control. By fabricating miniaturized systems with spatially controlled material properties, we propose a multiparametric platform to address key biological questions, including vascularization^45,49^; the influence of material properties on stem cell fate^2,4,50^; cell–matrix interactions,^51,52^ such as those between cancer-associated fibroblasts (CAFs) and dense matrices in cancer research^53^; and ECM homeostasis including matrix secretion and MMP-mediated degradation.^51^ These applications highlight the potential of our approach to advance both fundamental research and translational biomaterial design.

## Material and methods

### GelMA synthesis

Gelatin methacryloyl (GelMA) was synthesized following the one pot method developed by Shirahama *et al*. with some modifications.^54^ Gelatin (type A from porcine skin, bloom 300, Sigma Aldrich, G2500) was dissolved at 10% (w/v) in 0.25 M of carbonate-bicarbonate (CB) buffer solution (pH 9.4) for 1 h at 55°C under stirring. Methacrylic anhydride (MMA) (Sigma Aldrich, 276685) was added dropwise under vigorous agitation (500 rpm), while maintaining the pH at 9.4 using 5 M NaOH. The reaction proceeded at 55°C under constant stirring (500 rpm) for 1 h. Then the reaction was stopped by adding 100 mL of DI and adjusting the pH to 7.4. After being filtered (0.2 μm), the solution was dialyzed using dialysis tubing of 12-14 kDa of molecular weight cut-off (MWCO) (Spectra/Por® 4, Spectrumlabs) for 5 days against DI water at 40°C, replacing the water every 2–3 hours. Then the samples were frozen in liquid nitrogen for 20 min and lyophilized for one week at 0.02 mbar and –80°C using Lyoquest –85°C (Telstar). The resulting lyophilized GelMA powders were finally recovered and falcons were sealed with parafilm to prevent rehydration and stored at −20°C until use.

### Gel precursor solutions preparation

Stock solutions were prepared by dissolving all solid products in PBS at the following concentrations: 15% w/v GelMA, 10% w/v Dextran (Sigma Aldrich, 31392, 500 kDa), 2% w/v LAP (Lithium phenyl-2,4,6-trimethylbenzoylphosphinate, Sigma-Aldrich, 900889). Gel precursor solutions were obtained by mixing the stock solutions together with PBS to reach the desired concentrations of each component. The solutions were kept protected from light at 4°C for up to two weeks, and were heated to 37°C at least one hour before use. For sterile experiments, the gel precursor solution was filtered through 0.22 µm membrane before use. To allow visualization of the gel, methacrylated Rhodamine B (Polyscience, 23591) was dissolved in DI water to 1 mg/mL, filtered through 0.22 µm membrane, and added to the gel precursor at 1:100 volume ratio. The LAP concentration was kept at 0.25% w/v for all gel precursor solutions.

### Influence of GelMA/Dextran concentration

35 gel precursor solutions were prepared to span a concentration range from GelMA 3.5% to 5.5% w/v and Dextran 1.5% to 3.5% w/v. The solutions were heated to 37°C before being injected in a multi-channel microfluidic chip (ChipShop, Fluidic 561) which was placed in an incubator at 15°C for 10 minutes to allow phase separation before being crosslinked using a 405 nm UV curing LED (Amazon) with a power of 20 mW/cm^2^ for 1 min 30 (total dose 1.8 J/cm^2^). The chip was then observed using a confocal microscope (Leica SP8 with 40X water objective).

### Transmission electron microscopy (TEM)

Hydrogel samples were fixed with 2% glutaraldehyde in Sorensen buffer (0.1 M, pH = 7.4) for 1 h and washed with Sorensen phosphate buffer (0.1 M) for 12 h. They were then postfixed with 1% OsO 4 in Sorensen buffer (Sorensen phosphate 0.05 M, glucose 0.25 M, OsO 4 1%) for 1 h, washed twice with distilled water, and stained with 2% uranyl acetate aqueous solution for 12 h. Following the method described by Kiyama *et al*.^55^, water was gradually replaced with a water-ethanol solution. Specifically, the hydrogels were sequentially washed with H_2_O:EtOH mixtures in the ratios of 30:70, 15:85, 5:95, and finally 0:100%. This process was then repeated by using acetone. The samples were ultimately embedded in epoxy resin (Epon-Araldite) by following a stepwise protocol: acetone/epoxy resin mixtures of 2:1, 1:1, and 1:2, followed by pure epoxy resin, without any intermediate drying step. After 48 h of polymerization at 60°C, ultrathin sections (70 nm) were mounted on 100-mesh collodion-coated copper grids and poststained with 3% uranyl acetate in 50% ethanol (Delta Microscopies, Mauressac, France) and with 8.5% lead citrate before being examined using CCD detectors on the Hitachi electron microscope HT7700 at an accelerating voltage of 80 kV. All images were acquired at a magnification of x50,000 and saved in 16-bit format. Solid fraction was then estimated through image analysis on Fiji by applying flat field correction, automatic threshold, and by measuring the black to white pixel ratio (Figure S7).

### Rheometer measurements

Rheological measurements were carried out in oscillatory shear on a MCR 702e MultiDrive rheometer (Anton Paar, France) equipped with an active Peltier hood (temperature and atmosphere control) with a lower quartz glass plate. The 405 nm UV LED source (Omnicure LX500) was positioned under a quartz glass plate. Gelation temperature tests were performed to evaluate changes in the storage (G’) and loss modulus (G’’) associated with physical cross-linking in a soft gel containing 4.5% w/v GelMA (G4.5), a stiff gel containing 9% w/v GelMA (G9) and a LLPS-induced gel containing 4.5% w/v GelMA and 2.5% w/v dextran (G4.5D2.5), as well as to characterize the evolution of G’ and G’’ upon UV irradiation. For this purpose, 60 μL of GelMA or GelMA/dextran solution was placed between a quartz glass plate and an 8 mm diameter stainless steel plate (PP08/S) with a 1 mm gap. The samples were subjected to a constant shear strain of 1% and a constant frequency of 1 Hz while the temperature was gradually increased or decreased according to a predefined profile. The experimental workflow consisted of six steps: 1) the temperature was maintained at 37°C for 1 min; 2) the temperature was decreased from 37 to 15°C at a constant rate of 6.3°C/min over 3.5 min; 3) the temperature was maintained at 15°C for 30 min; 4) UV irradiation was applied at 15°C for 3 min; 5) the temperature was increased from 15 to 37°C at a constant rate of 11°C/min over 2 min; 6) the temperature was maintained at 37°C for 5 min.

### Influence of temperature

A custom temperature controller was fabricated to allow temperature monitoring while imaging under microscope. Peltier elements (TEC1-12705) were used as heating and cooling elements to finely monitor the temperature of a copper plate on which was placed the sample. The controller is based on Arduino using the PID library for precise feedback control. Device wiring diagram and code are provided in Supplementary Information 1. The gel precursor was pipetted between two glass coverslips with a 250 µm thick PDMS spacer ring. The temperature was monitored in real time and the sample was observed through phase-contrast microscopy.

### Pore size characterization

The pore size was determined with Fiji on confocal and phase contrast images. For the confocal images, images were binarized by thresholding, smoothed using a median filter, and the pore size distribution was determined using the local thickness function. For the phase contrast images, Fast Fourier Transform (FFT) analysis was performed to determine the characteristic lengths of periodic elements. The main peak of the FFT radial profile was found to match the pore size measured on confocal images. The image analysis procedure is detailed in Figure S3.

### Thermal camera imaging

Thermal images were taken using a FLIR (X6580SC) camera. The coverslip to be imaged was coated with high emissivity black paint to reduce infrared light reflection during imaging.

### DMD patterning

The gel precursor solution was injected between two coverslips with a 250 µm thick PDMS spacer at 37°C, and the LLPS was initiated by cooling down either using an incubator or directly under the microscope using the Peltier device. Photopatterning was carried out under microscope using a DMD-based photopatterning equipment (Primo, Alvéole) with 405 nm light and an exposure of dose of 7 mJ/mm^2^, unless otherwise mentioned.

### Cell culture

Human Lung Fibroblasts (ATCC, PCS-201-013) were cultivated using conventional protocols in DMEM GlutaMAX (Gibco 10566016) supplemented with 10% fetal bovine serum (FBS) and 1% penicillin/streptomycin. Cells were split when confluent using trypsin and kept until P6. Before seeding in the gel, cells were starved by incubating them in medium without FBS overnight to reset fibroblast activity.

### Cell encapsulation in the porous gel

Cells were trypsinized, washed once with PBS at 37°C and resuspended in the gel precursor solution at 37°C at a concentration of 700k cells/mL. PDMS microfluidic chips with 4 parallel channels (each being 18 mm long, 3 mm wide, and 350 µm thick) were used to cast the gel on methacrylated glass coverslips. The PDMS chips and coverslips were sterilized in 70% ethanol for 30 min and dried in an oven at 60°C for 1 hour prior to cell seeding. The gel precursor containing cells was then thoroughly introduced in the channels (13 µL), before being submitted to phase separation for 8 min, either in a cooling incubator or using the Peltier device. The chips were then exposed with 405 nm light for 40s at 20 mW/cm2. The chips were placed at 37°C for 3 min to dissolve non-crosslinked components, and the PDMS chip was separated from the coverslip. The coverslip with gels were then cultured in static conditions in petri dishes filled with cell culture medium for up to 15 days, with a medium change every 3 to 4 days.

### Methacrylated coverslip preparation

Glass coverslips (24 x 33 mm) were cleaned by wiping with soap and paper, then immersed for 5 min in a Piranha solution (1 volume of H_2_SO_4_ for 1 volume of 30% H_2_O_2_). After thorough rising with DI water, the slides were dried and immersed in a solution of ethanol containing 2% v/v MAPTMS (Trimethoxy[3-(methylamino)propyl]silane) and 0.01% v/v acetic acid for 1h. Slides were then washed with ethanol, dried, and heated on a hotplate at 110°C for 30min to complete the reaction.

### PDMS microfluidic chip fabrication

The PDMS chip was fabricated using soft lithography. A silicon wafer with positive SU-8 patterns was fabricated using conventional lithography, rendered hydrophobic by silanization using 1H,1H,2H,2H-Perfluorooctyltriethoxysilane vapors, and used as a mold for PDMS casting (Sylgard 184, 1:10 crosslinking agent to polymer ratio). The channels dimensions were 18×3 mm in x-y and 150 µm deep. After curing overnight at 60°C, the PDMS was peeled off from the mold, cut into indivial chips and the inlets were manually punched.

### Cell staining and imaging

The gels containing cells were fixed using 4% paraformaldehyde in PBS for 30 min, permeabilized using 0.1% Triton X-100 in PBS for 15 min, blocked in 5% FBS in PBS for 2h, and incubated 1h with 1:400 Phalloidin Alexa Fluor 488, 2 µg/mL DAPI and 1% FBS in PBS. Three PBS washing steps with 5 min incubation were performed between each step. The cells were observed using epifluorescence (Zeiss Axio Observer) and confocal (Leica SP8 with 40X water objective; and Revvity Opera Phenix with 20X water objective) microscopes. For confocal imaging, a drop of SlowFade Glass mounting medium (ThermoFisher) was added onto the gels to improve imaging depth.

### Cell image processing

The quantification was performed using a custom macro on Fiji on the epifluorescence images of gels fixed at day 1, 3, 7, 11 and 15. The image was first split into 30, 600 µm wide, sections from left to right with increasing porosity On each section, the pore size was analyzed using the FFT method (Figure S3) on the bright field channel, and the nuclei were counted on the DAPI channel using the Find Maxima function. The macro returned a table with the data for each section including x-position within the gel, pore size, and nuclei density. All samples were performed in triplicates, and the error bars show the standard deviation between samples.

## Supporting information

Supporting Information 2: Arduino code

## Acknowledgements

This work has received the support of Toulouse Initiative for Research’s Impact in Society (TIRIS) ANR-22-EXES-0015. This work has benefited from a government grant managed by the Agence Nationale de la Recherche under the France 2030 program, reference ANR-24-EXME-0002 (PEPR MEDOoC). This work has received the support of Natural Sciences and Engineering Research Council (NSERC) Vanier Graduate Scholarship and NSERC Discovery Grant (RGPIN 2025-05408). This work was partly supported by the LAAS-CNRS micro and nanotechnologies platform, a member of the French Renatech network and by the LAAS-CNRS characterization platform. This work was partly supported as part of the MultiFAB project funded by FEDER European Regional Funds and French Région Occitanie (16007407/MP0011594). The authors also thank Astrid Canivet and David Sagnat from the Organoid facility of IRSD (Toulouse).

## Authors contributions

SD, DGM, JF, JMM and LM led and designed the project. SD, DGM, APO, DFA, CB and JF designed experiments. SD, DGM and MP analyzed data. SD, DGM, APO, DFA, LB, AA, CB, DG, RP, MH and JF carried out experiments. SD and LM wrote the manuscript.

## References

1. Puiggalí□Jou, A., Hui, I. B., Fernandez□Rico, C. & Zenobi□Wong, M. The Space Within: How Architected Voids Promote Tissue Formation. Advanced Materials e07385 (2025) doi:10.1002/adma.202507385.

2. Han, S., Kim, J., Lee, G. & Kim, D. Mechanical Properties of Materials for Stem Cell Differentiation. Adv. Biosys. 4, 2000247 (2020).

3. Wu, Y. et al. Viscoelastic extracellular matrix enhances epigenetic remodeling and cellular plasticity. Nat Commun 16, 4054 (2025).

4. Engler, A. J., Sen, S., Sweeney, H. L. & Discher, D. E. Matrix Elasticity Directs Stem Cell Lineage Specification. Cell 126, 677–689 (2006).

5. Roloson, E. B. et al. Collagen Scaffold Viscoelasticity Regulates Muscle Cell Phenotype. Adv Healthcare Materials e02775 (2025) doi:10.1002/adhm.202502775.

6. Polz, M. et al. Cellular Responses to Mechanical Cues Across Scales: From Fundamental Insights to Translational Potential. Adv Healthcare Materials 15, e71400 (2026).

7. Drury, J. L. & Mooney, D. J. Hydrogels for tissue engineering: scaffold design variables and applications. Biomaterials 24, 4337–4351 (2003).

8. Rezakhani, L. et al. Recent advances in hydrogels applications for tissue engineering and clinical trials. Regenerative Therapy 26, 635–645 (2024).

9. Xiao, S. et al. Gelatin Methacrylate (GelMA)-Based Hydrogels for Cell Transplantation: an Effective Strategy for Tissue Engineering. Stem Cell Rev and Rep 15, 664–679 (2019).

10. Aigoin, J. et al. Comparative Analysis of Electron Microscopy Techniques for Hydrogel Microarchitecture Characterization: SEM, Cryo-SEM, ESEM, and TEM. ACS Omega 10, 14687–14698 (2025).

11. Annabi, N. et al. Controlling the Porosity and Microarchitecture of Hydrogels for Tissue Engineering. Tissue Engineering Part B: Reviews 16, 371–383 (2010).

12. Maksoud, F. J. et al. Porous biomaterials for tissue engineering: a review. J. Mater. Chem. B 10, 8111–8165 (2022).

13. Annabi, N. et al. Controlling the Porosity and Microarchitecture of Hydrogels for Tissue Engineering. Tissue Engineering Part B: Reviews 16, 371–383 (2010).

14. Grenier, J. et al. Mechanisms of pore formation in hydrogel scaffolds textured by freeze-drying. Acta Biomaterialia 94, 195–203 (2019).

15. Savina, I. N., Ingavle, G. C., Cundy, A. B. & Mikhalovsky, S. V. A simple method for the production of large volume 3D macroporous hydrogels for advanced biotechnological, medical and environmental applications. Sci Rep 6, 21154 (2016).

16. Gun’ko, V. M., Savina, I. N. & Mikhalovsky, S. V. Cryogels: Morphological, structural and adsorption characterisation. Advances in Colloid and Interface Science 187–188, 1–46 (2013).

17. Park, E., Ryu, J. H., Lee, D. & Lee, H. Freeze–Thawing-Induced Macroporous Catechol Hydrogels with Shape Recovery and Sponge-like Properties. ACS Biomater. Sci. Eng. 7, 4318–4329 (2021).

18. Joukhdar, H. et al. Ice Templating Soft Matter: Fundamental Principles and Fabrication Approaches to Tailor Pore Structure and Morphology and Their Biomedical Applications. Advanced Materials 33, 2100091 (2021).

19. Djemaa, I. B., Auguste, S., Drenckhan-Andreatta, W. & Andrieux, S. Hydrogel foams from liquid foam templates: Properties and optimisation. Advances in Colloid and Interface Science 294, 102478 (2021).

20. Tang, Y. et al. In situ gas foaming based on magnesium particle degradation: A novel approach to fabricate injectable macroporous hydrogels. Biomaterials 232, 119727 (2020).

21. Yi, S. et al. Micropore□Forming Gelatin Methacryloyl (GelMA) Bioink Toolbox 2.0: Designable Tunability and Adaptability for 3D Bioprinting Applications. Small 18, 2106357 (2022).

22. Sergeeva, A., Feoktistova, N., Prokopovic, V., Gorin, D. & Volodkin, D. Design of Porous Alginate Hydrogels by Sacrificial CaCO_3_ Templates: Pore Formation Mechanism. Adv Materials Inter 2, 1500386 (2015).

23. Han, L.-H., Lai, J. H., Yu, S. & Yang, F. Dynamic tissue engineering scaffolds with stimuli-responsive macroporosity formation. Biomaterials 34, 4251–4258 (2013).

24. De Rutte, J. M., Koh, J. & Di Carlo, D. Scalable High□Throughput Production of Modular Microgels for In Situ Assembly of Microporous Tissue Scaffolds. Adv Funct Materials 29, 1900071 (2019).

25. Escudero, M. et al. Scalable Generation of Pre□Vascularized and Functional Human Beige Adipose Organoids. Advanced Science 10, 2301499 (2023).

26. Deo, K. A. et al. Granular Biphasic Colloidal Hydrogels for 3D Bioprinting. Adv Healthcare Materials 13, 2303810 (2024).

27. Daly, A. C. Granular Hydrogels in Biofabrication: Recent Advances and Future Perspectives. Adv Healthcare Materials 13, 2301388 (2024).

28. Ribeiro, L. S., Gaspar, V. M., Sobreiro□Almeida, R., Camargo, E. R. & Mano, J. F. Programmable Granular Hydrogel Inks for 3D Bioprinting Applications. Adv Materials Technologies 8, 2300209 (2023).

29. Sheikhi, A. et al. Microfluidic-enabled bottom-up hydrogels from annealable naturally-derived protein microbeads. Biomaterials 192, 560–568 (2019).

30. Matsunaga, Y. T., Morimoto, Y. & Takeuchi, S. Molding Cell Beads for Rapid Construction of Macroscopic 3D Tissue Architecture. Advanced Materials 23, (2011).

31. Xu, F., Dodd, M., Sheardown, H. & Hoare, T. Single-Step Reactive Electrospinning of Cell-Loaded Nanofibrous Scaffolds as Ready-to-Use Tissue Patches. Biomacromolecules 19, 4182–4192 (2018).

32. Song, J., Michas, C., Chen, C. S., White, A. E. & Grinstaff, M. W. From Simple to Architecturally Complex Hydrogel Scaffolds for Cell and Tissue Engineering Applications: Opportunities Presented by Two□Photon Polymerization. Adv Healthcare Materials 9, 1901217 (2020).

33. Mukasheva, F., et al. Design and characterization of 3D printed pore gradient hydrogel scaffold for bone tissue engineering. Bioprinting 39, e00341 (2024).

34. Picado-Tejero, D., Mendoza-Cerezo, L., Rodríguez-Rego, J. M., Carrasco-Amador, J. P. & Marcos-Romero, A. C. Recent Advances in 3D Bioprinting of Porous Scaffolds for Tissue Engineering: A Narrative and Critical Review. JFB 16, 328 (2025).

35. Arizpe Tafoya, A. V., et al. Multi-material 3D Bioprinting of Complex Heterogeneous Constructs for In-vitro Mechanobiology Studies. Preprint at 10.26434/chemrxiv.15007731/v1 (2026).

36. González□Martínez, D. A., et al. A Systematic Study of GelMA□Carbopol Bioinks for High□Fidelity Extrusion 3D Bioprinting at Physiological Temperatures. Adv Healthcare Materials e71248 (2026) doi:10.1002/adhm.71248.

37. He, Z. & He, S. Two-photon polymerization of hydrogel cellular scaffolds. Optics Communications 574, 131161 (2025).

38. Van Der Sanden, B., Gredy, L., Wion, D. & Stephan, O. 3D two-photon polymerization of smart cell gelatin – collagen matrixes with incorporated ruthenium complexes for the monitoring of local oxygen tensions. Acta Biomaterialia 130, 172–182 (2021).

39. Mu, J. et al. Phase-separated hydrogels for advanced biomedical engineering: From material design to applications. Materials Today Bio 38, 103096 (2026).

40. Halwes, M. et al. Tuneable Hydrogel Porosity via Dynamic Tailoring of Spinodal Decomposition. Advanced Science 12, e04265 (2025).

41. Hori, A. et al. One-Step Formation of Microporous Hydrogel Sponges Encapsulating Living Cells by Utilizing Bicontinuous Dispersion of Aqueous Polymer Solutions. ACS Appl. Bio Mater. 2, 2237–2245 (2019).

42. Ben Messaoud, G., Aveic, S., Wachendoerfer, M., Fischer, H. & Richtering, W. 3D Printable Gelatin Methacryloyl (GelMA)□Dextran Aqueous Two□Phase System with Tunable Pores Structure and Size Enables Physiological Behavior of Embedded Cells In Vitro. Small 19, 2208089 (2023).

43. Stefanopoulou, E., Ben Messaoud, G., Salazar Ortiz, R., Fischer, H. & Richtering, W. Controlled in situ acidification enables the 3D printability of GelMA–dextran aqueous two-phase hydrogel with unique interconnected porosity. RSC Appl. Polym. 4, 308–319 (2026).

44. Dudaryeva, O. Y. et al. Tunable Bicontinuous Macroporous Cell Culture Scaffolds via Kinetically Controlled Phase Separation. Advanced Materials 37, 2410452 (2025).

45. Dudaryeva, O. Y. et al. Multi□scale Engineered Vasculature and Hierarchical Porosity via Volumetric Bioprinting□Guided Photopolymerization□Induced Phase Separation. Advanced Materials 38, e21171 (2026).

46. Müller, M. Z. et al. Cell-guiding microporous hydrogels by photopolymerization-induced phase separation. Nat Commun 16, 4923 (2025).

47. Gao, D. et al. Macroporous scaffolds based on biomass polymers and their applications in wound healing. Journal of Bioresources and Bioproducts 10, 14–31 (2025).

48. Hinton, T. J. et al. Three-dimensional printing of complex biological structures by freeform reversible embedding of suspended hydrogels. Sci. Adv. 1, e1500758 (2015).

49. Wang, Y. et al. Mechanical strategies to promote vascularization for tissue engineering and regenerative medicine. Burns & Trauma 12, tkae039 (2024).

50. Olivares-Navarrete, R. et al. Substrate Stiffness Controls Osteoblastic and Chondrocytic Differentiation of Mesenchymal Stem Cells without Exogenous Stimuli. PLoS ONE 12, e0170312 (2017).

51. Narasimhan, B. N. & Fraley, S. I. Matrix degradation enhances stress relaxation, regulating cell adhesion and spreading. Proc. Natl. Acad. Sci. U.S.A. 122, e2416771122 (2025).

52. Naba, A. Mechanisms of assembly and remodelling of the extracellular matrix. Nat Rev Mol Cell Biol 25, 865–885 (2024).

53. Najafi, M., Farhood, B. & Mortezaee, K. Extracellular matrix (ECM) stiffness and degradation as cancer drivers. J of Cellular Biochemistry 120, 2782–2790 (2019).

54. Shirahama, H., Lee, B. H., Tan, L. P. & Cho, N.-J. Precise Tuning of Facile One-Pot Gelatin Methacryloyl (GelMA) Synthesis. Sci Rep 6, 31036 (2016).

55. Kiyama, R. et al. Nanoscale TEM Imaging of Hydrogel Network Architecture. Advanced Materials 35, 2208902 (2023).

