## Supporting Information for "Interconnected Macroporous GelMA Scaffolds for Cell Culture through Temperature Controlled Spinodal Decomposition"

### Supporting Figures

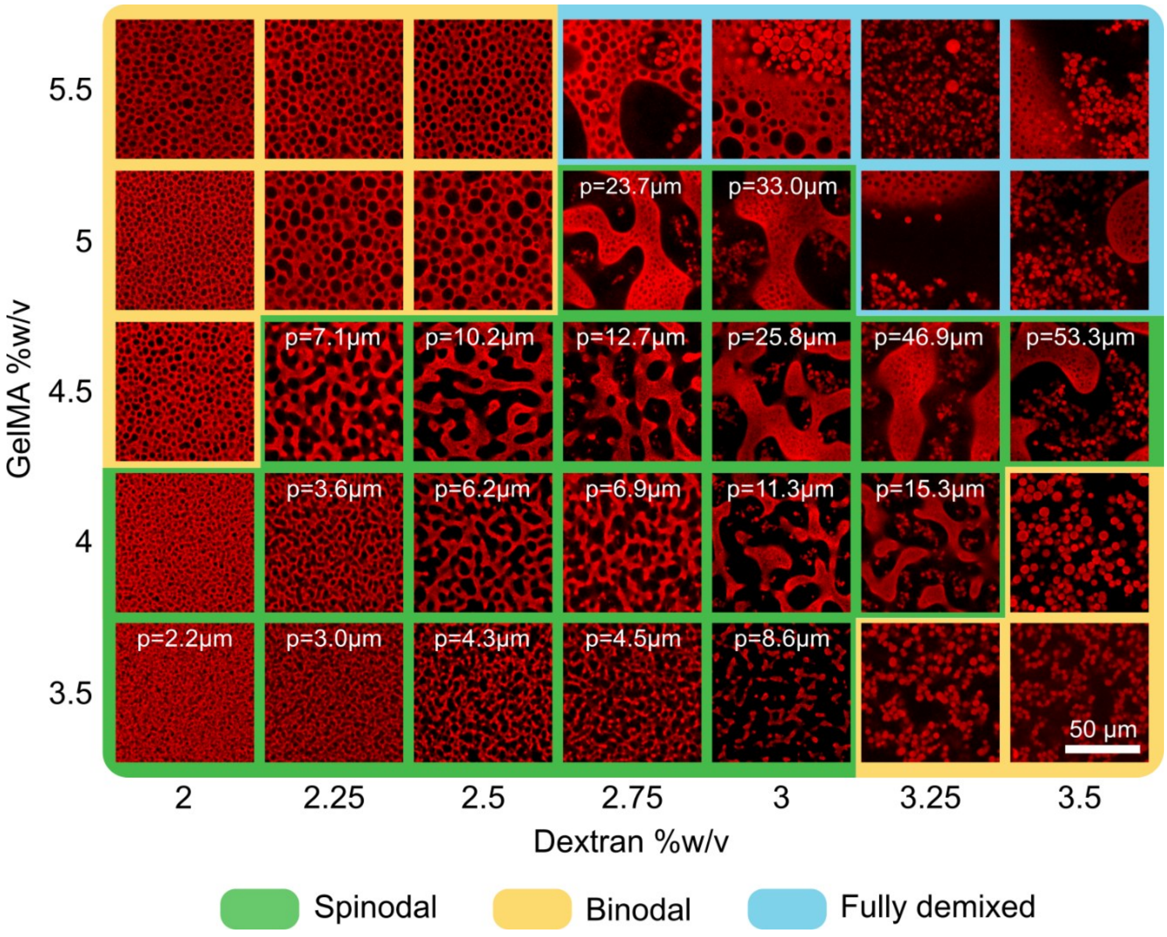


***Figure S1:*** *Microstructures obtained with different gel compositions at 15°C. p gives the average pore size.*

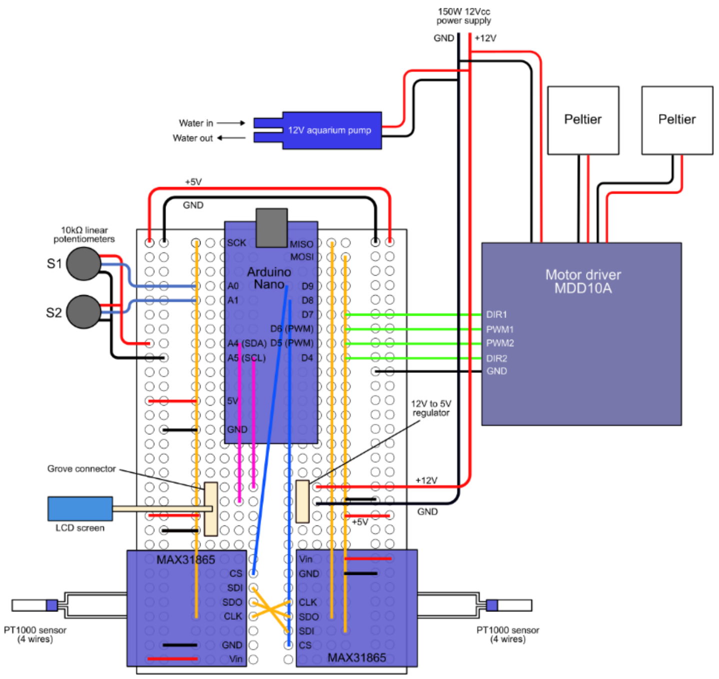


***Figure S2:*** *Wiring diagram of the Peltier controller device (for 1 or two Peltier elements). Supply list and Arduino program are provided in Supporting Information 1 and 2*

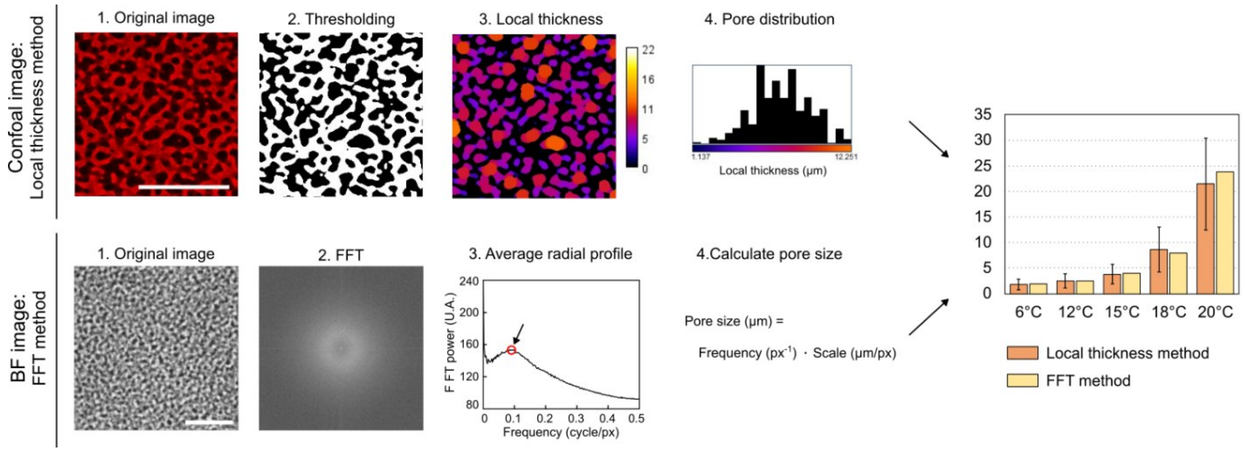


***Figure S3:*** *Image processing for pore size characterization. The first method employs confocal microscopy images as the starting material. It involves thresholding followed by local thickness analysis to generate a porosity distribution. Error bars represent the standard deviation of pore distribution, measured in the local thickness histogram. The second method, designed for bright-field images, utilizes a Fast Fourier Transform (FFT)-based approach to extract the dominant feature sizes. In this method, the FFT is first computed, and the average radial profile is extracted. This profile represents the power spectrum of the FFT, with frequency (x-axis, in cycles per pixel) being inversely proportional to the feature size (in micrometers). By detecting the peak in this profile, the main feature size can be calculated. Our results demonstrate that both methods yield comparable values, confirming that the detected feature size corresponds to the actual pore size distribution.*

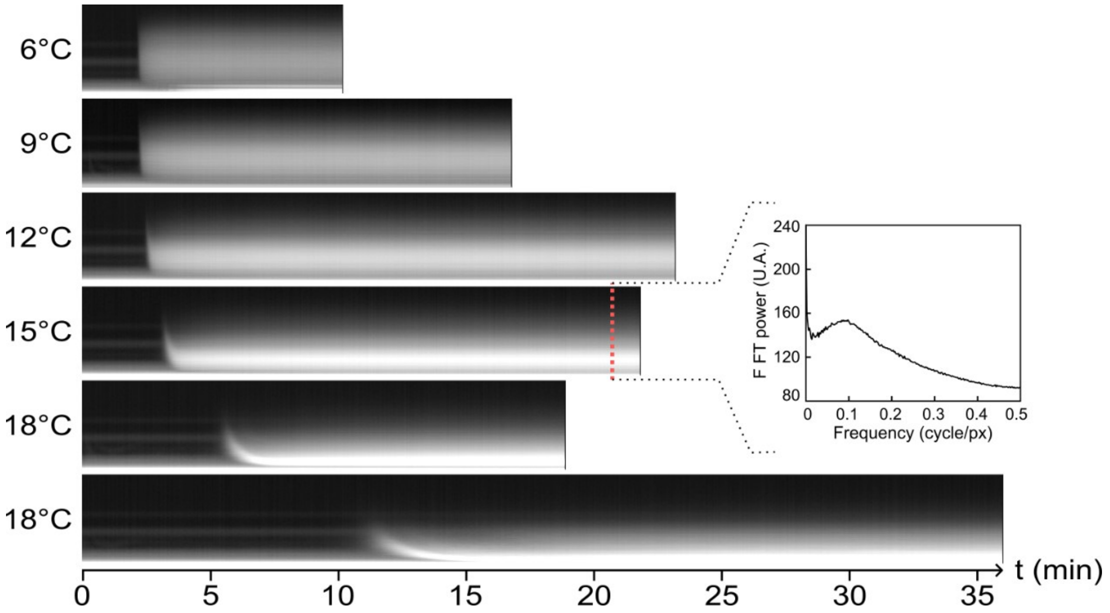


***Figure S4:*** *Kymographs of the FFT profile extracted from videos of LLPS at different temperatures (Video S2). On each kymograph, the x-axis corresponds to time, and vertical lines are power specturm of the FFT radial profile.*

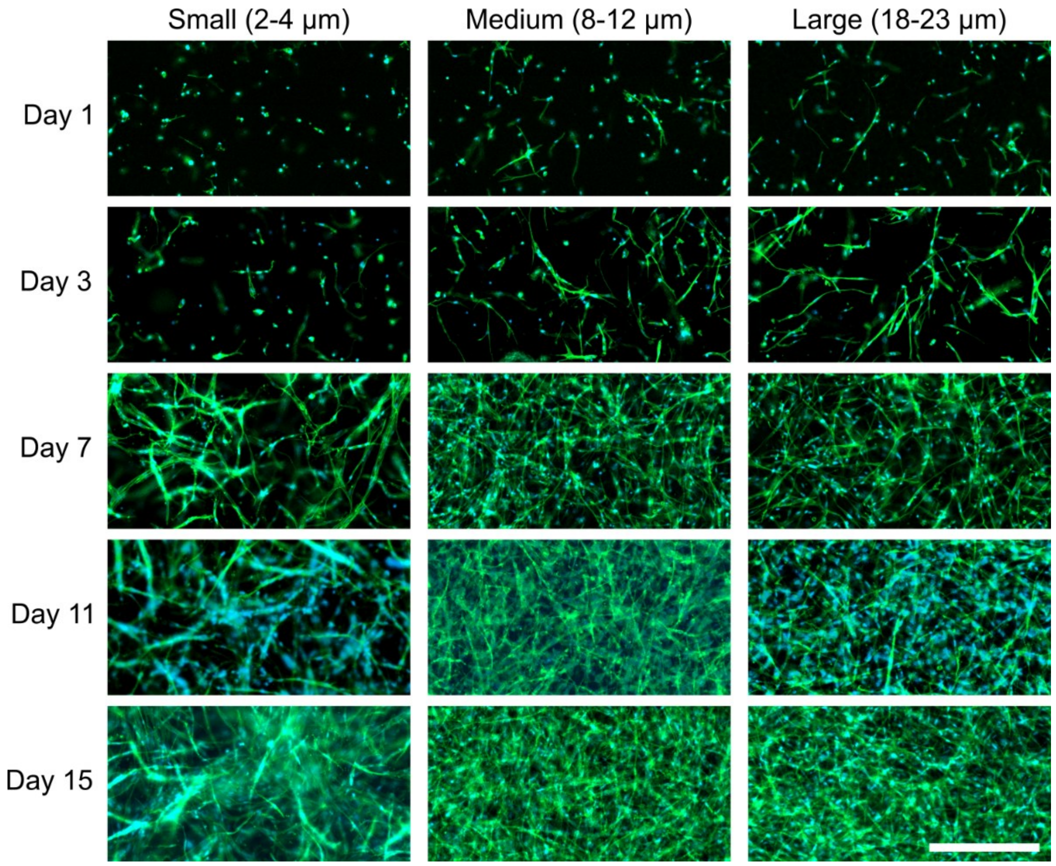


***Figure S5:*** *Epifluorescence micrographs of cells growing in different porosity regions over 15 days. Scale bar 500 µm.*

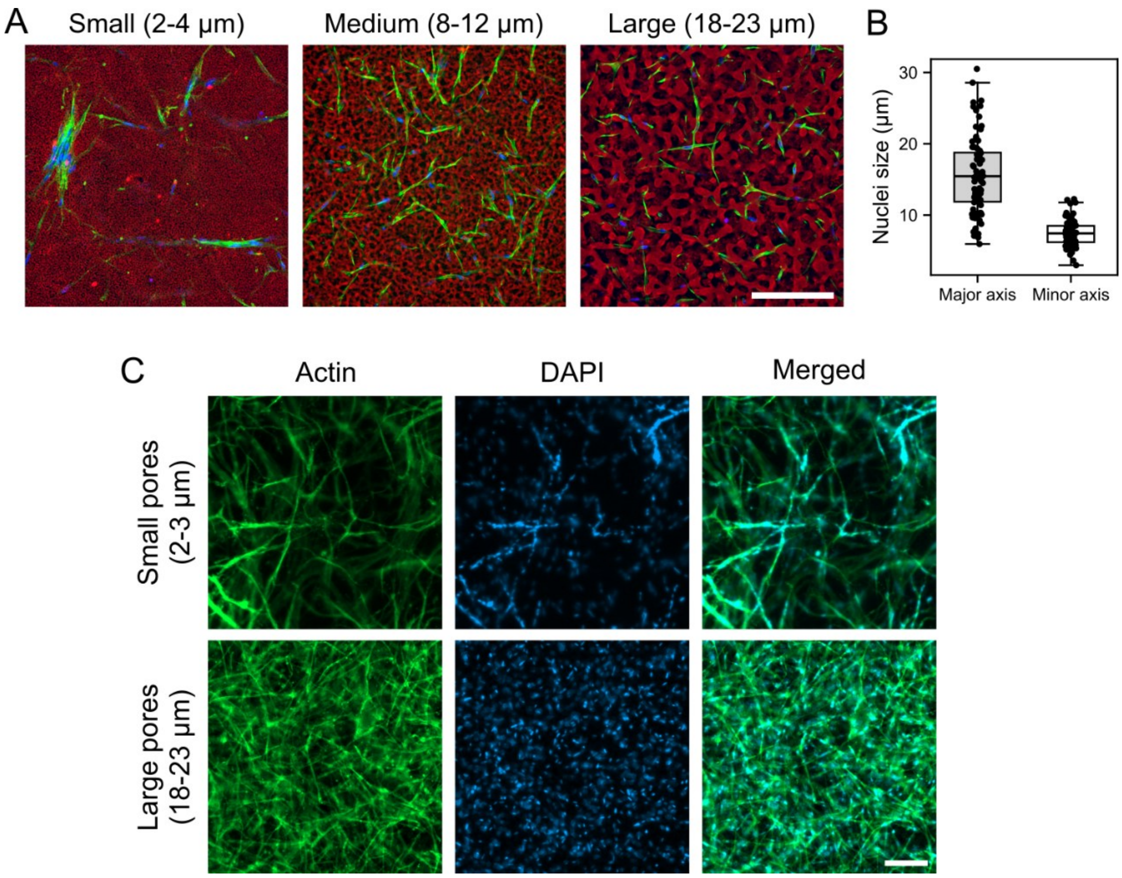


***Figure S6:*** *Morphology change in response to cell confinement.* ***A.*** *Confocal images at day 15 on three regions of different porosities. This highlights the mesenchymal/collective to amoeboid transition by increasing the porosity, as well as the maximum growth in medium (8-12 µm) porosity.* ***B.*** *Nuclei size measured on the 3 pictures in A: Mean minor axis = 7.5 µm, Mean major axis = 15.6 µm. (n = 78 nuclei)* ***C.*** *Epifluorescence images of cells in small (2-3 µm) and large (18-23 µm) porosity gel at day 11. Nuclei are evenly distributed in large pores regions, while they align into bundles in confined regions. Scale bar 200 µm.*

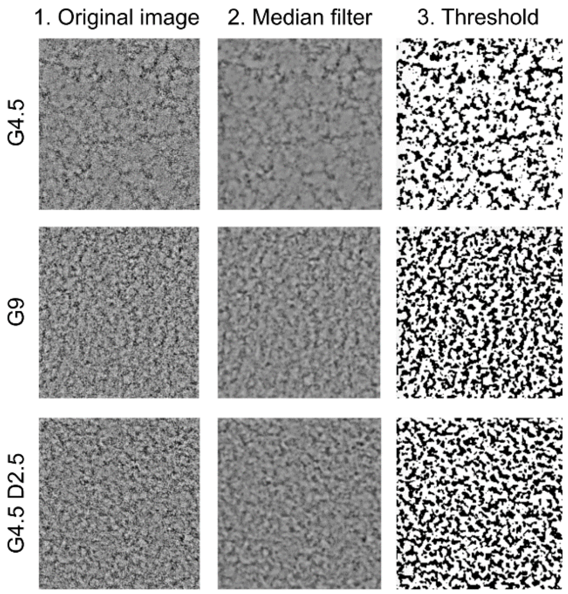


***Figure S7:*** *TEM images analysis. Solid fractions are calculated by measuring the black pixel area over the white on the thresholded TEM pictures. G4.5 $\phi$=0.29, G9% $\phi$=0.40, G4.5D2.5 $\phi$=0.39*

**Supporting Information 1: Peltier temperature controller**

Supply list:

- TEC1-12705 Peltier modules (x2)
- Water cooling block 40x40mm, Amazon (x2)
- Copper plates, 60x40x1 mm (x2)
- Aquarium pump (x1)
- Sapphire window, Edmund Optics 75-142 (x1)
- EC360 thermal glue (x1) used for thermal contact between Peltiers, cooling blocks, copper plates and temperature sensors.
- Prototyping board, uPesy medium board (x1)
- Arduino nano (x1)
- MDD10A 2x10A DC motor controller (x1)
- PT1000 temperature sensor, RS PRO 293-8446 (x2)
- MAX31865 temperature sensor interface (x2)
- 10kΩ linear potentiometer (x2)
- I2C compatible LCD screen for Arduino, Grove 104020113 (x1)
- 12VCC power supply 150W (x1)
- S13V10F5 voltage regulator 12 to 5VCC (x1)

**Supporting Information 2: Arduino code**

File attached

**Supporting videos**

**Video S1:** Spinodal decomposition video

**Video S2:** Spinodal decomposition at different temperatures

**Video S3:** Photopatterning

**Video S4:** Confocal stack
